# Near-atomic resolution Cryo-EM structure of Mayaro virus identifies key structural determinants of alphavirus particle formation

**DOI:** 10.1101/2021.01.14.425226

**Authors:** David Chmielewski, Jason Kaelber, Jing Jin, Scott C. Weaver, Albert J. Auguste, Wah Chiu

## Abstract

Mayaro virus (MAYV) is an arthritis-inducing alphavirus circulating in the Americas, with potential to rapidly emerge in new geographical regions and populated environments. Intraparticle heterogeneity has typically limited atomic resolution structures of alphavirus virions, while imposing icosahedral symmetry in data processing prevents characterization of non-icosahedral features. Here, we report a near-atomic resolution cryo-EM structure of the MAYV E1-E2-E3-CP subunit by addressing deviations from icosahedral symmetry within each virus particle. We identified amino acid contacts at E1 protein interfaces forming the icosahedral lattice and investigated their effect on MAYV growth through site-directed mutagenesis. Further, mutation of a short stretch of conserved residues in E2 subdomain D, near an unidentified “pocket factor” including E2Y358, significantly reduced MAYV growth and provides strong evidence that this unknown factor influences assembly. Further, a symmetry-free reconstruction revealed the MAYV virion is not strictly icosahedral, suggesting defects in global symmetry may be a feature of the virus particle budding process. Our study provides insights into alphavirus assembly and suggests a common path in the formation of spherical, enveloped viruses, leading to particle imperfections.

## Introduction

The *Alphavirus* genus (family *Togaviridae)*, contains arthropod-transmitted pathogens responsible for near-global epidemics in humans and livestock(Schmaljohn and McClain 2011; Strauss and Strauss 1994). Included in the genus are notable pathogens Chikungunya (CHIKV), Mayaro (MAYV) and Ross River virus (RRV), that account for millions of annual cases of debilitating, persistent polyarthritis across Africa, Asia, Australia, Europe and the Americas(Powers et al. 2001). These viruses continue to expand their geographic distribution, and the recent emergence of CHIKV in Asia and the Americas clearly demonstrates our inability to rapidly respond to and control their emergence(Kraemer et al. 2015; Tsetsarkin et al. 2007, 2009; Schuffenecker et al. 2006). MAYV primarily circulates within a sylvatic cycle in tropical forested regions of the Americas, though recent evidence of laboratory transmission by *Ae. aegypti, Ae. albopictus* and *Anopheline* vectors suggests potential adaptation to urban and peridomestic transmission(Acosta-Ampudia et al. 2018; Kantor et al. 2019; Torres et al. 2004; Mavian et al. 2017; Hotez and Murray 2017). MAYV exhibits significant cross-reactivity with other alphavirus species within the Semliki forest virus serogroup, and as a result cases are likely systematically misdiagnosed and underreported (Webb et al. 2019). Despite the increasing frequency of epidemics and expanding geographic range of many alphaviruses, there are no licensed vaccines or antiviral therapies for alphavirus infection.

The alphavirus genome consists of a single-stranded, ∼11.5kb (+)RNA genome that encodes six structural and four non-structural proteins. The non-structural proteins (nsPs1-4) are essential for genome replication and immune evasion. The structural polypeptide (capsid(CP)-E3-E2-6K/TF-E1) is produced from a sub-genomic promoter and cleaved both co- and post-translationally. CP is first auto-proteolytically processed from the structural polypeptide and specifically interacts with the MAYV genomic RNA (gRNA) to form nucleocapsids (NCs). E1 protein has membrane fusion activity while E2 interacts with cell surface receptors and mediates cellular entry via clathrin-mediated endocytosis (Lescar et al. 2001; Smith et al. 1995; W. Zhang et al. 2005; Rong Zhang et al. 2018). E3 is essential for proper folding of p62 (the precursor to E2) and provides stability of the E2-E1 heterodimer complex following cleavage by furin-like proteases during passage through the trans-Golgi network(Carleton et al. 1997; Heidner, Knott, and Johnston 1996; Mulvey and Brown 1995). 6K and TF are small lipophilic membrane proteins with largely unknown function, though 6K has been implicated in membrane ion permeability and virus budding(Melton et al. 2002; Firth et al. 2008; Lusa, Garoff, and Liljeström 1991; Loewy et al. 1995).

Previous cryo-EM studies of several alphaviruses have revealed the organization of the virus particle and individual E1-E2-(E3)-CP subunit(Paredes et al. 1993; Cheng et al. 1995; Rui Zhang et al. 2011). Particles are ∼70 nm in diameter, with 80 protruding spikes composed of trimerized E1-E2 heterodimer (20 icosahedral 3-fold and 60 quasi-3-fold spikes) arranged in *T=4* icosahedral symmetry. 240 copies of E1-E2 are embedded in the lipid membrane, with a direct interaction between the cytosolic tail of E2 and CP c-terminal binding-pocket(Taylor, Hanson, and Kielian 2007; Suomalainen, Liljeström, and Garoff 1992). Alphavirus particles bud from the plasma membrane, where E1-E2 complexes presented at the cell surface successively enwrap cytosolic NCs enclosing the (+)ssRNA genome. The mechanism of membrane scission required to release the nascent virion remains poorly understood, though it has been demonstrated the process is independent of the host ESCRT system(Taylor, Hanson, and Kielian 2007). While E2-CP interaction is essential to particle assembly, pre-formation of cytosolic NCs through CP-CP contacts is not a strict prerequisite to particle formation(Y. Zheng and Kielian 2015; Forsell et al. 2000).

Assembly of viral particles is a dynamic process, the endpoint of which depends greatly on the kinetics and thermodynamics of subunit association as well as fluctuations to the assembly conditions(Perlmutter and Hagan 2015). Production of virus particles with significant defects, such as missing capsomers or local deviations from the global symmetry, have been proposed to arise from independent assembly trajectories that never reach the globally-symmetric conformation of lowest energy (Perlmutter and Hagan 2015; Hagan and Chandler 2006). Rather than representing a population of unproductive nascent particles, virions with deviations from global symmetry have been proposed to possess a functional advantage due to the potential for more rapid disassembly and release of viral genome(Wang, Mukhopadhyay, and Zlotnick 2018). Traditional structural studies of spherical viruses by cryo-electron microscopy (cryo-EM) have relied on high degrees of global symmetry in the data processing steps to greatly increase averaging power and overcome the low signal of each radiation-sensitive virus particle image. Potential defects in each virus particle image are either masked by the symmetry applied in 3d-reconstruction, giving the appearance of a perfect particle, or computationally discarded by only selecting the most homogenous, ordered particles for the final structure.

Biochemical and 2D structural data indicate that purified CPs of Ross River virus do not form complete icosahedral shells *in vitro* and incorporate shell defects (Wang et al. 2015). To assess whether alphavirus particles possess defects in the icosahedral envelope and/or NC lattices, cryo-EM images of individual MAYV particles were first examined in 2D for potential defects. Due to the prevalence of deviations from global symmetry within each viral particle, an additional data processing protocol was developed at the sub-particle level to identify and discard those distorted regions of each virion image from the final average. This data processing increased the resolvability of a well-ordered sub-particle region, without imposing icosahedral symmetry in the reconstruction. Further, to better understand the deviations from global icosahedral symmetry among MAYV particles, we determined an asymmetric 3d-reconstruction of the virion. The resulting structure gives a novel insight into the organization of the envelope glycoprotein (GP) and NC layers, with relevance to the poorly understood mechanism of alphavirus budding.

By accounting for imperfections in the alphavirus particle lattice, we report a near-atomic resolution cryo-EM structure of MAYV and the corresponding all-atom model, and identify sites relevant to virus assembly at side-chain resolution. To validate the importance of specific interfacial amino acid interactions, we perform structure-guided mutagenesis experiments and assess the effects on MAYV replication. Based on these results, we identify a novel site of lateral envelope interactions at the interface of neighboring trimers that likely influences assembly of the MAYV icosahedron, and validate a separate target for disruption of a lipophilic pocket with relevance to E1-E2 heterodimer assembly.

## Results

### MAYV particles possess defects in the icosahedral lattice

The cryo-EM structure of the MAYV virion was first solved to ∼4.7Å by imposing icosahedral symmetry, revealing a structural conservation typical of the alphavirus family (Figure 1a-d). However, visual inspection of individual 2D images of MAYV particles revealed significant heterogeneity, with virions apparently pleomorphic in shape (Figure 2). Further inspection of the particle images revealed a few apparent abnormalities: namely (1) missing capsomere units, (2) distorted and/or extended side of particle and (3) multi-cored particles containing multiple, typically two, NCs enwrapped in a lipid envelope decorated with envelope GPs.

**Figure 1.**
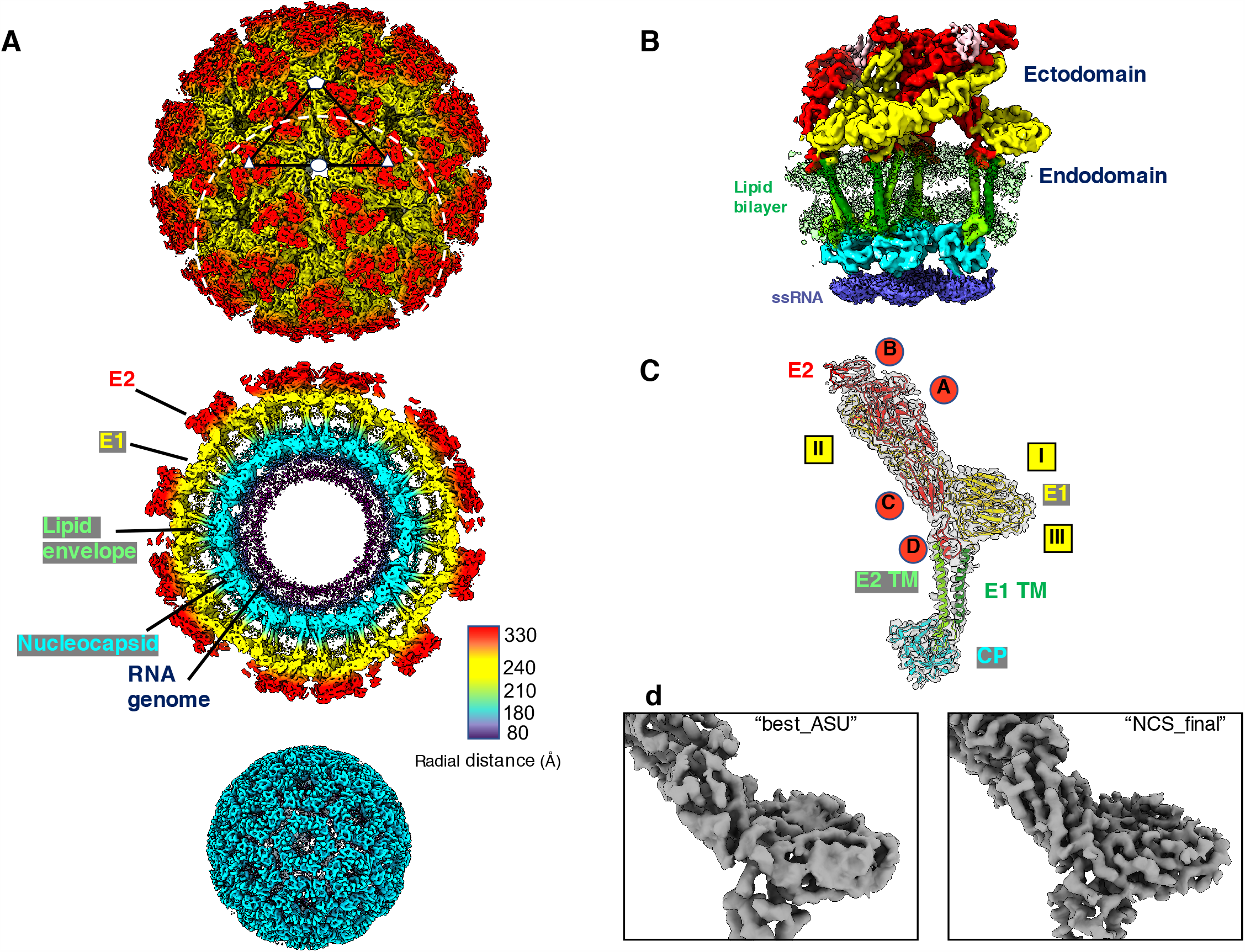
Cryo-EM structure of MAYV. (a) Radially-colored surface representation of MAYV, with a central section to reveal interior of particle. Protein, lipid and nucleic acid features are labeled in the central section surface view. Symmetry axes of virus particles represented by white shapes (pentagon:5-fold, triangle:3-fold, ellipse:2-fold. Masked region used for focused-refinement protocol of subparticles depicted as white circles with dashed lines. Nucleocapsid density extracted and colored as cyan. (b) Side-view representation of MAYV asymmetric unit (ASU) consisting of four E1(yellow)-E2(red)-E3(pink)-CP(cyan) subunits arranged as a quasi-3-fold trimer and a single subunit contributing to an icosahedral-3-fold trimer. (c) Density of E1-E2-CP subunit after NCS-averaging of four quasi-equivalent subunits within one ASU, with fit backbone model of each protein. Domains of E2 ectodomain (red circles) and E1 ectodomain (yellow squares) labeled. (d) Representative density of E1/E2 ectodomain after focused-refinement of subparticles (“best_ASU”) and NCS averaging (“NCS_final”).

**Figure 2.**
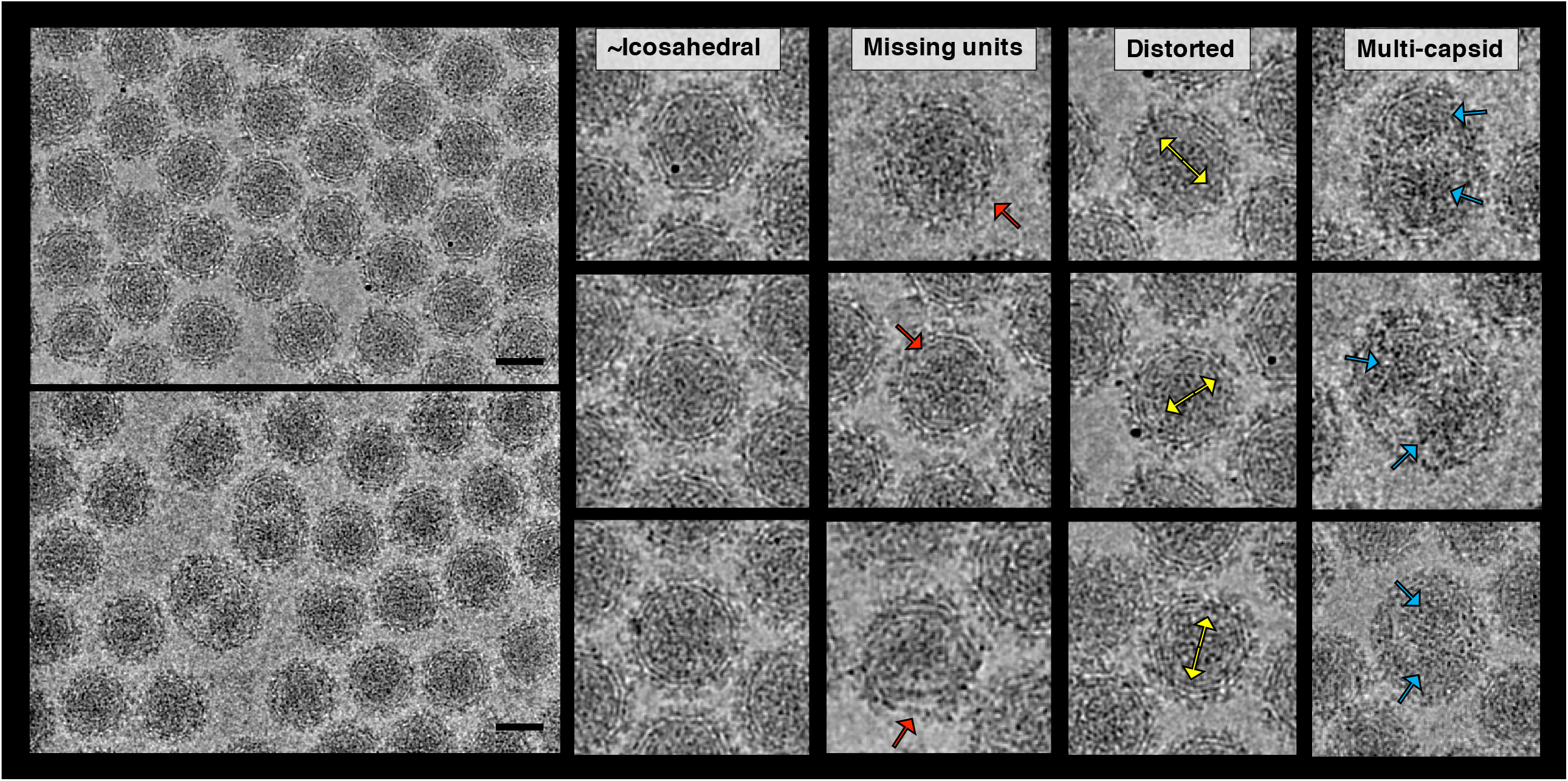
Representative images of purified MAYV particles display significant heterogeneity. Selected images are of particles belonging to each of four characteristic virus morphologies (“icosahedral”, “missing units”, “distorted”, and “multi-capsid”). Red arrows represent locations of apparently missing or disrupted glycoprotein and/or nucleocapsid layers, yellow arrows represent apparent axis of elliptical distortion, and blue arrows highlight location of multiple capsids within one enveloped MAYV particle. Scale bar: 50 nm.

To address the effect of incorrect assignment of asymmetric unit (ASU) orientations throughout each virus particle image containing defects from global symmetry, an additional data processing protocol was implemented. First, 60 masked subparticles, corresponding to the 60 ASUs of the virus, were generated from each raw particle image based on the rough icosahedral orientations. Each masked subparticle, centered on a unique ASU and including additional padding density (Figure 1a), was then refined against the MAYV penton reference with local c5 symmetry and 3d-orientation search constraints. Each ASU subparticle was scored by phase residual agreement to reference, regardless of the virus particle which it originated, and the highest-scoring subset was retained for the final average. The additional protocol improved the map resolution ∼0.3Å beyond the icosahedrally-averaged map to 4.4Å, allowing for the first slight separation of beta-strands in the E1-E2 ectodomain and observation of pitch in the transmembrane (TM) helices (Figure EV 1a-b).

Non-crystallographic symmetry (NCS) was used to average the four quasi-equivalent E1-E2-E3-CP subunits within the MAYV ASU, further improving the definition of the peptide backbone and side chains (Figure EV 1b). The final NCS-averaged subunit density map of the envelope was determined to 4.2Å resolution and C-terminal protease domain of CP to 4.4Å resolution by gold-standard (0.143) FSC. Subsequent Q-score analysis of the agreement between model and density suggests the E1-E2 ectodomain structure meets the criteria of a map far better than 4.2Å resolution because many side chain densities are apparent and rotamers can be properly modeled (Figure EV 2a-c) (Pintilie et al. 2020a). Comparison of the MAYV atomic model with those of other alphaviruses revealed high similarity between the protein domains (Figure EV 4). E1 ectodomain is divided into three domains: I, II (contains fusion loop) and III (lies parallel to lipid envelope) (Figure 1c, Movie S1). E2 ectodomain is divided into four domains: A (putative receptor binding function), B (putative receptor binding function, covering fusion loop), C and a β-ribbon connector region (Figure 1c). Q-score analysis of the CP supports the lower measured resolution in this region of the map, as the average residue Q-score (Qresidue) of 0.48 is lower (corresponding to a lower resolution map) than that of the E1 ectodomain (Qresidue 0.57) and E2 ectodomain (Qresidue 0.54) (Figure EV 3a-c). Density corresponding to two N-linked GlcNac sugar moieties were observed in the subunit density map, one at the surface-exposed residue E1 N141 and the second at E2 N362 along the E1-E2 interface (Figure EV 5a-b). We also observe density of cleaved E3 in our map in association with E2 beta-ribbon connector, in agreement with its positioning in other alphavirus cryo-EM maps (Figure 1b).

### MAYV particles have structurally ordered and disordered poles

To test the global symmetry of particles, an asymmetric reconstruction of MAYV was determined by refining the orientation of each particle to one unique solution instead of imposing icosahedral symmetry constraints. This symmetry-relaxation resulted in a converged map of MAYV at ∼9Å resolution. In contrast to the seemingly “perfect” MAYV virion map calculated with icosahedral symmetry, the symmetry-free map contains deviations from global symmetry. A highly structured side of the symmetry-free reconstruction, which we termed this side the “leading” pole, is opposed by the most disordered region on the other side of the virion, we refer to this as the “trailing” pole (Figure 3a-d). The leading pole, approximately one-third of the particle, exhibits ordered GPs, trans-envelope helices, and internal CPs of the nucleocapsid layer consistent with the icosahedral reconstruction. Density becomes progressively weaker starting just above the midpoint of the virion and ending at the disordered trailing pole. The density of GPs, TM helices and NC at the trailing pole is largely missing, and appears distorted, suggesting this region contains significant deviations from icosahedral symmetry.

**Figure 3.**
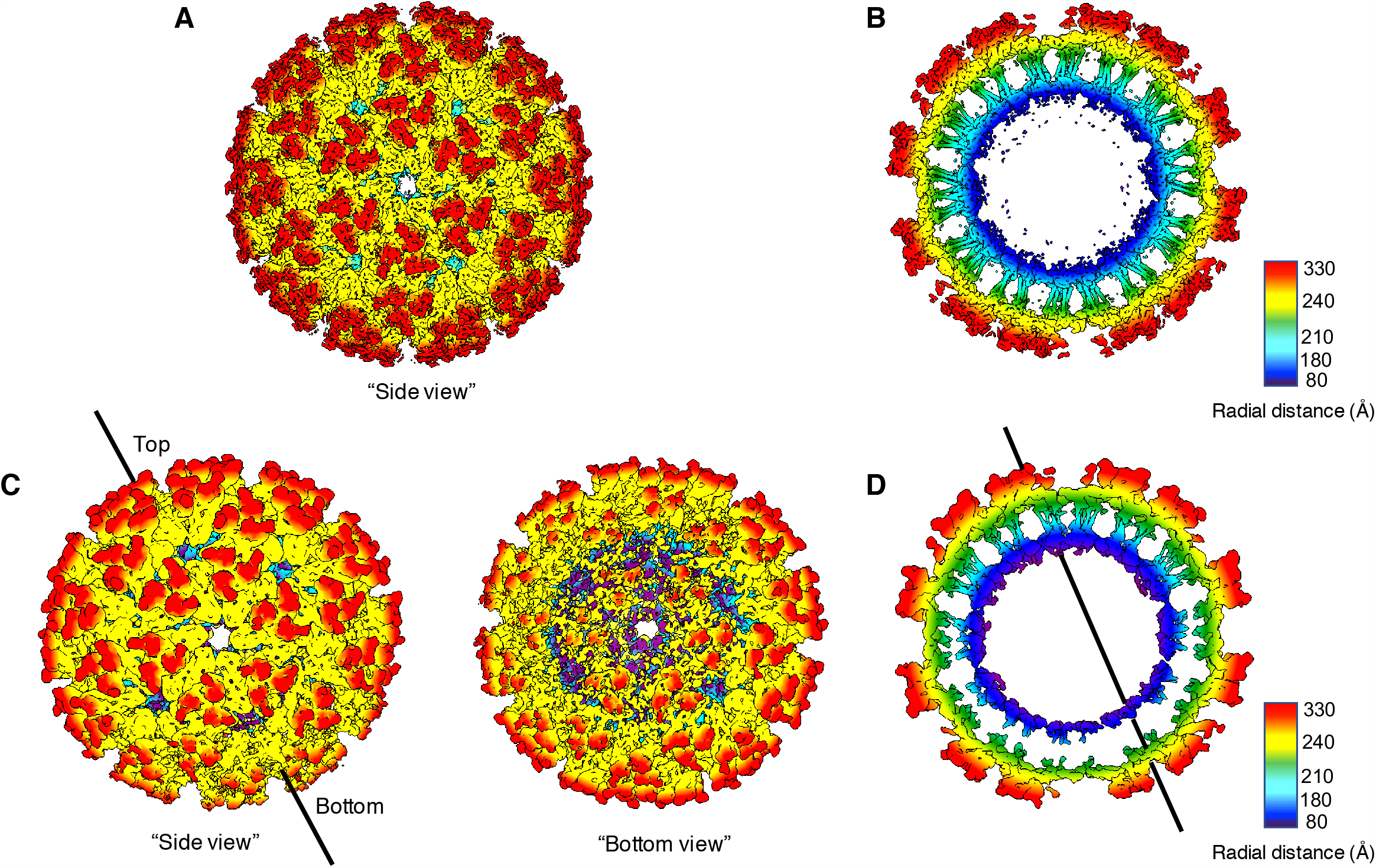
Cryo-EM reconstructions of MAYV with and without icosahedral symmetry. (a) Surface view of MAYV reconstruction with icosahedral symmetry imposed during data processing with radial surface coloring. (b) Central section through the icosahedral reconstruction reveals consistent structure throughout the virus particle. (c) Surface view of asymmetric MAYV reconstruction without global symmetry imposed during refinement, colored radially. Side view displays ordered icosahedral density while bottom view displays weaker icosahedral features. Black line illustrates the axis of most ordered density directly opposing the particle surface with least ordered density. (d) Central section through the asymmetric reconstruction shows a clear polarity to order of the glycoprotein ectodomains (yellow/red), lipid envelope/glycoprotein helices (green/cyan) and nucleocapsid (dark blue/purple).

After observing the polarity in icosahedral order in the asymmetric reconstruction, we sought to better understand how the nucleocapsid was positioned relative to the GP layer. We found that the NC density in the asymmetric reconstruction is positioned equidistant from the GP layers at both the leading and trailing poles. Interestingly, the NC region of most disorder at the trailing end correlates with the region of most disordered GPs, lacking density of the TM helices and showing weak, distorted density for the GP trimers. This suggests that NCs are not completely icosahedral in the mature MAYV virion, and CP organization is likely modulated by the organization of the external GP layer through direct contacts.

### Lateral envelope interactions on the virus surface

80 trimers composed of E1-E2 heterodimers occupy 60 quasi-three-fold (q3) and 20 icosahedral three-fold (i3) positions of the MAYV *T=4* icosahedral lattice. These trimers are arranged as 12 pentons with five-fold symmetry and 20 hexons with a two-fold symmetry axis (Figure 4a). The lateral protein-protein interactions of the MAYV GPs can be understood by observing the two distinct interfaces between neighboring trimers: those formed by (1) i3-q3 trimers (“type I”) and (2) q3-q3 trimers (“type II”). The type I trimer interface is extensive (∼990Å^2^) and relatively flat, with five evenly-distributed regions of inter-trimer contacts with quasi-2-fold symmetry (Figure 4b). In comparison, the type II interface (∼790Å^2^) is situated at a greater angle, with three regions of contacts positioned near the five-fold axis.

**Figure 4.**
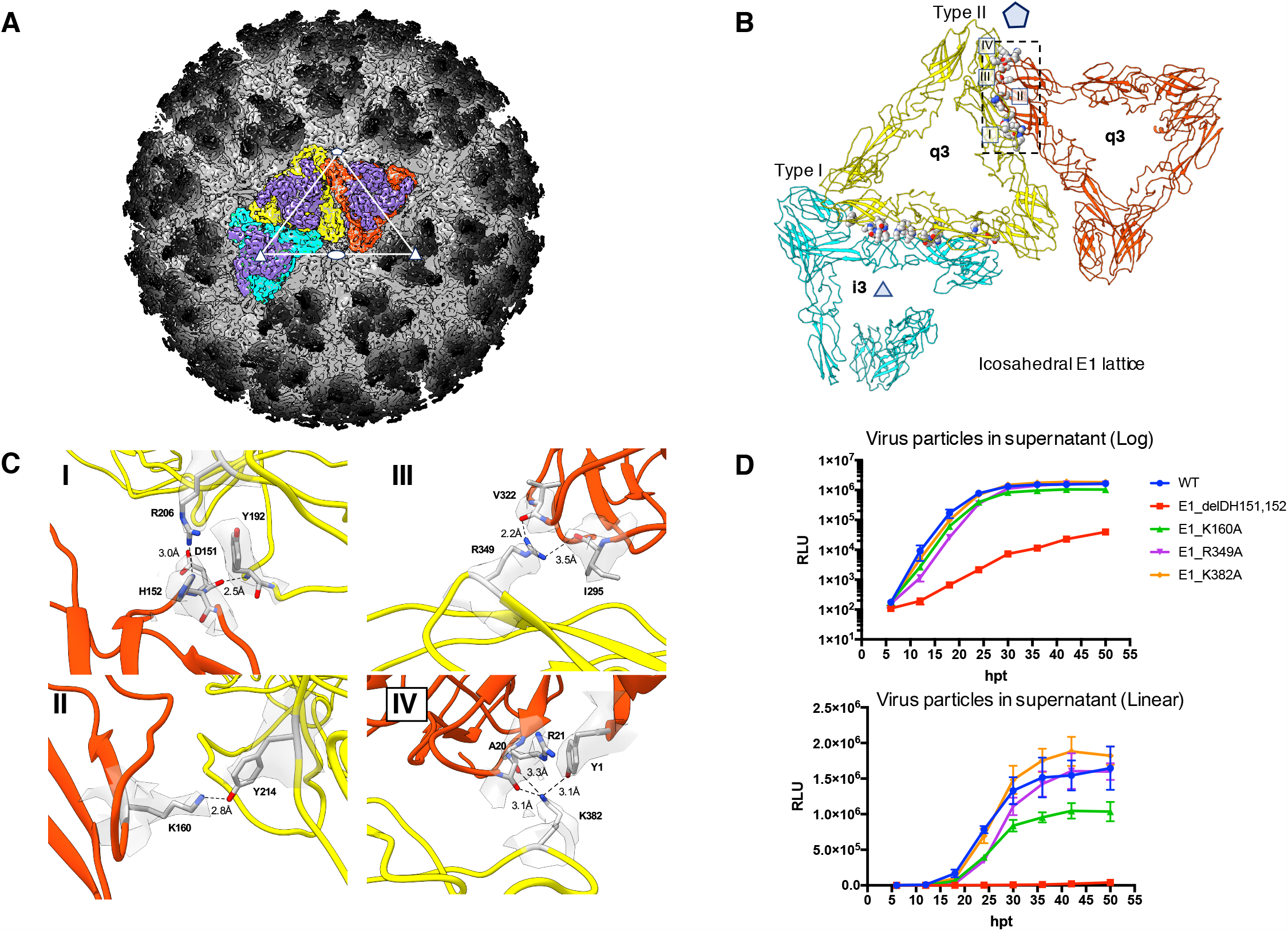
Lateral interactions between MAYV surface spikes. (a) Radially colored surface representation of MAYV, with three neighboring surface trimers colored (E2-purple, E1-cyan,yellow,orange). White triangle shows the position of 5-fold axis (pentagon), 3-fold axes (triangles) and 2-fold axis (ellipse). (b) Molecular interactions across E1 molecules of neighboring GP trimers displayed as grey spheres. Type I interactions between quasi-3fold (yellow) and icosahedral-3fold trimers (cyan) and Type II interaction between two quasi-3fold trimers (yellow and orange). The Pentagon represents the position of the 5-fold axis. (c) Atomic interactions identified at the Type II interfaces (I-IV), with zoned density around interacting residues. (d) Growth curves of MAYV infectious reporter clone (WT and E1 mutants), measured by relative light units (RLU) at time points up to 48hpi and displayed in linear and log scales.

Side-chain interactions at Type II inter-trimer interfaces were identified with PyMol and PDBePISA tool, and validated by assessing agreement between model and density using a “Q-score” protocol (Figure EV 2b)(Pintilie et al. 2020b). Type II contacts were investigated for influence on the icosahedral lattice due to the more limited overall interface area and location near the five-fold axis. We generated MAYV-NLuc reporter virus, with E2 N-terminus nanoluciferase fusion, to quantitatively measure viral titers of WT and several MAYV point mutants and a deletion targeting the identified inter-trimer contacts. MAYV-NLuc genomic RNAs (gRNAs) were transcribed and capped *in vitro* and transfected into BHK cells via electroporation. Growth kinetics were assessed by measuring luciferase activities in the culture supernatant every six hours for 48 hours post-transfection (hpt) (Figure 4d).

Within the type II inter-trimer interface, a significant contact region is formed between E1 domain I (E1DI) and E1DII of the neighboring trimer (Figure 4c). E1D151 main-chain carbonyl (loop DIG_0_) forms a H-bond with main-chain amino group of Y192 of neighboring E1, while E1H152 can form a polar interaction with R206 of the neighboring E1 molecule. Deletion construct E1ΔD151H152, made to abolish the inter-trimer H-bond formed between peptide backbones, showed a significant 2-3 log reduction in growth relative to WT at 18hpt and 2-log reduction at 48hpt. Nearby, E1K160 (loop DIH_0_) forms a H-bond with E1Y214 of the neighboring q3 trimer. Mutant E1K160A reduced virus assembly relative to WT approximately 2-fold at measured time points. This difference is clearly presented in linear scale (Figure 4d).

Additional type II contact regions near the five-fold axis were identified across E1DIII of two E1 molecules from neighboring quasi-3-fold trimer. The first involves E1K382 that forms multiple H-bonds with main-chain carbonyls of A20/R21 and Y1 of the neighboring E1 (Figure 4c). Mutant K382A had no effect on virus assembly. The other interaction, located directly at the five-fold axis, involves H-bonds between E1R349 and main-chain carbonyls of V322 and I295. Mutant E1R349A showed 2-fold reduction in growth at 24 hpt, after which virus production gradually approached that of WT. Interestingly, in our structure I295 side chain atoms do not make inter-trimer contacts as proposed for S295 in CHIKV and SFV, though a polar interaction does exist nearby between E1T305 and E1S369 of the neighboring q3 trimer.

### Unassigned factor in lipophilic pocket near viral membrane

Cryo-EM maps of alphavirus particles have consistently revealed density corresponding to the major structural proteins E1-E2-CP, and in some cases the cleaved E3 (Rui Zhang et al. 2011; Garoff, Simons, and Renkonen 1974; Basore et al. 2019). As first noted in the map of Venezuelan equine encephalitis virus (VEEV), additional unassigned density exists in a hydrophobic pocket formed by E2 subdomain D (subD) and the top of E1 transmembrane helix closest to the outer leaflet of the lipid envelope(Rui Zhang et al. 2011). Recently, this density has been termed a “pocket factor” and its identity suggested to be the hydrophobic tail of a phospholipid(Chen et al. 2018). We observe the unassigned factor in our map of MAYV (Figure 5a) in the highly lipophilic pocket (Figure 5c, Appendix Movie S1), where it runs roughly parallel to E2subD and contacts residues E2H362, E2Y358 and E2P351. Interestingly, E2Y358 is completely conserved in all mosquito-borne alphaviruses while E2H362 is highly conserved (Figure 5b, Appendix Figure S1). In our map, the pocket factor is well-resolved at density thresholds where E2 domain B is not, suggesting the factor is more stably positioned than the flexible distal tip of E2. In addition to contacts between subD and the lipophilic factor, we observe the H-bond between E2H348 and E1T403, first noted in the structure of VEEV (residues E2H348, E1S403), that is proposed to stabilize the orientation of E2 tail and E1 ectodomain lattice (Rui Zhang et al. 2011). Consistent with the recent SINV cryo-EM structure, E2H352 does not form any E1-E2 contacts in our map as previously suggested (Byrd and Kielian 2017), instead E2P351 forms a contact with E1W407 (Figure 5c).

**Figure 5.**
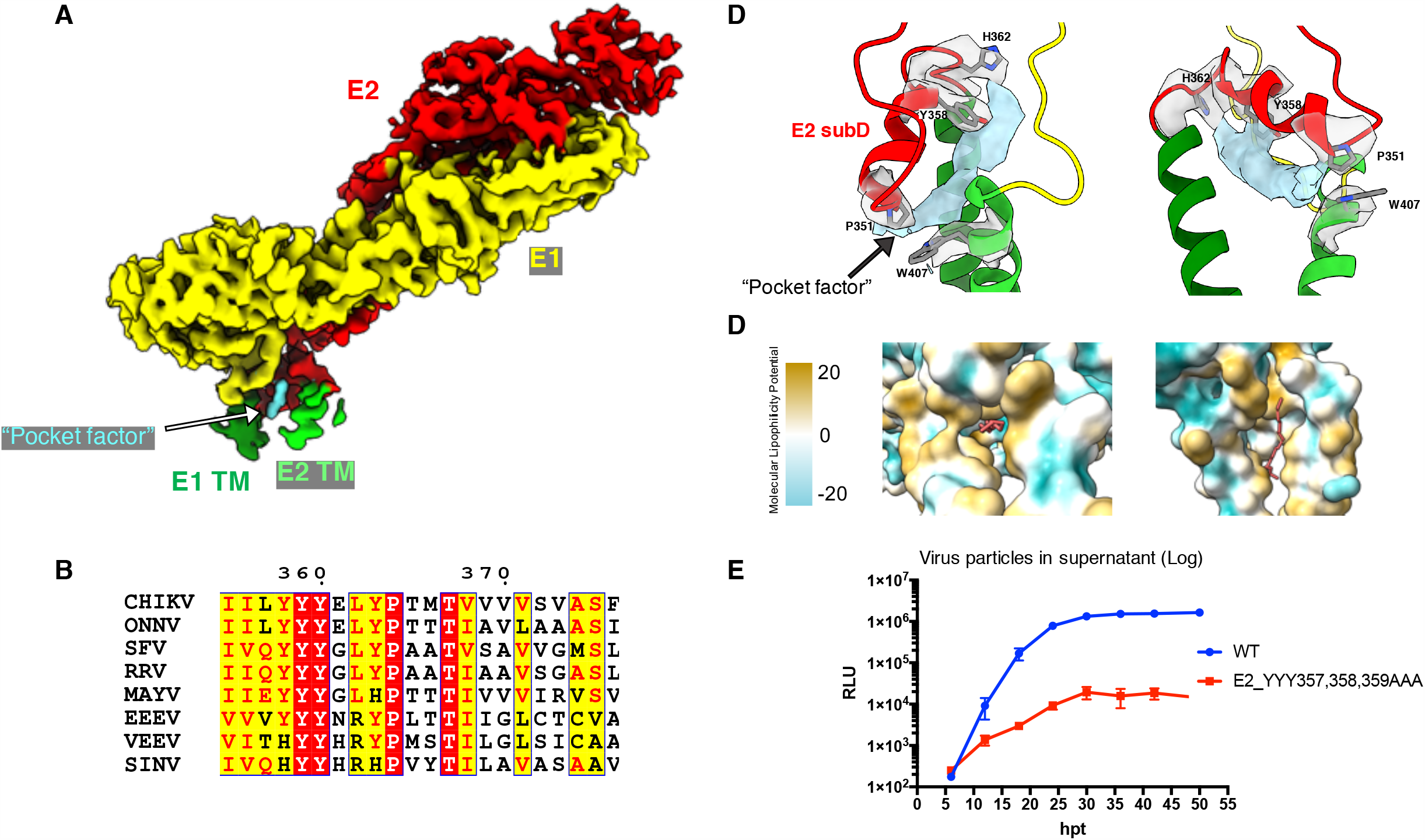
Unassigned density in the cryo-EM structure of the MAYV E1/E2 heterodimer. (a) E1-E2 heterodimer density (E1-yellow, E2-red) with unassigned “pocket factor” density (sky blue). (b) E2 sequence alignment of multiple alphaviruses shows complete conservation of E2Y358 and E2Y359 (MAYV numbering). Highly conserved residues shown with red background and white font, somewhat conserved residues shown with yellow background and red font. Sequence alignment performed using NPS and ClustalW, formatted with ESPript. (CHIKV-Chikungunya virus, ONNV-O’nyong-nyong virus, SFV-Semliki Forest virus, RRV-Ross River virus, EEEV-Eastern equine encephalitis virus, VEEV-Venezuelan encephalitis virus, SINV-Sindbis virus). (c) Zoom-in view of E2subD with zoned density around E2 residues forming contacts (H362,Y358,P351) with the pocket factor. E1 residue W407 forms a E1-E2 contact with P351. (d) Pocket factor and surrounding density as represented by molecular lipophilicity potential, a three-dimensional representation of lipophilicity. Hydrophobic tail of a phospholipid, as previously proposed (PDB:6IMM), is rigidly fit into the pocket factor density and displayed (salmon). (e) Growth curves of MAYV infectious reporter clone after transfection, including WT and E2 Y357A,Y358A,Y359A triple-alanine mutant as measured by RLU.

To assess the importance of highly conserved E2subD tyrosines to virus assembly and/or replication, a triple substitution of Y357,Y358,Y359 with alanines was introduced into our MAYV-NLuc reporter virus. In all cryo-EM structures with suitable resolution, conserved residue E2Y358 (numbering differs) contacts the central region of pocket factor (Figure 5c). Virus assembly, monitored as described previously, was significantly reduced for the triple mutant E2_YYY357,358,359AAA by 2-3 log at measured time points (Figure 5e). Loss of important contacts between E2subD and the pocket factor could potentially destabilize the highly lipophilic pocket in its pre-fusion conformation, leading to instability of the E1-E2 heterodimer assembly.

## Discussion

The classical model of virus assembly requires each subunit to be arranged in equivalent (*T=1*) or quasi-equivalent (*T>1*) chemical environments on the viral capsid(Crick and Watson 1956; Caspar and Klug 1962). The association of individual viral subunits into icosahedrons can be envisioned for ideal cases where kinetic or thermodynamic traps are avoided, but the concurrent processes of subunit assembly and budding of a planar membrane presents additional challenges(Hagan and Chandler 2006). Molecular simulation suggests steric clashes between GP trimers and extreme negative membrane curvature of the late-stage budding “neck” can prevent incorporation of GPs at the trailing end of the alphavirus particle (Lázaro, Mukhopadhyay, and Hagan 2018a). Our visual inspection of MAYV particles and focused classification of ASUs revealed significant defects to global icosahedral symmetry within purified virus particles. While we cannot rule out mechanical damage during virus purification or vitrification, we envision a cellular origin to these deviations. Our observation of multi-cored particles, where GP trimers directly contact only one side of the NCs, suggests an icosahedral GP layer completely wrapping around a NC is not required for particle release. These unique particles support a model where virions spontaneously achieve membrane scission or unknown host factors facilitate release without completion of the icosahedron at the trailing end. It is also possible that these multi-cored particles relieve kinetic stalls of the budding shells, proposed to occur at mid- to late-stages of budding, through lateral GP self-interactions (Lázaro, Mukhopadhyay, and Hagan 2018b).

In our asymmetric MAYV structure, we observed polarity in the resolvability of both envelope GP trimers and NC capsomers from the ordered, leading pole to a more disordered, trailing pole. To achieve this average, an icosahedral structure at one side is opposed by either missing capsomers or ASUs in non-icosahedral positions. Interestingly, asymmetric cryo-EM reconstructions of an immature and mature flavivirus revealed similar polarity of structural order, with more extreme differences between the poles and NC positioning in the immature particle containing GP trimers (Therkelsen et al. 2018). Both alphaviruses and flaviviruses require GPs and NC for virus budding, though the exact contribution of each component during assembly is not well understood. The similarity between symmetry-free flavivirus and alphavirus reconstructions can potentially indicate a shared mechanism of GP-driven assembly for these icosahedral, enveloped virus families. The ordered, leading pole suggests lateral GP trimer interactions can drive initial formation of the icosahedral lattice, while the apparent deviations to icosahedral symmetry at the “trailing” end are expected due to steric constraints at late stages of budding. In the alphavirus family, NCs are more icosahedral in virions, presumably through direct interactions with symmetrically-organized GPs. Our observations that CP density is lower resolution than the GP ectodomain in the E1-E2-E3-CP subunit map, and that the NC does not show clear icosahedral density at the “trailing” end of the asymmetric reconstruction, add evidence that it is the GP layer that imparts icosahedral symmetry to the NC through direct E2-CP interactions (Figure EV 3d). This agrees with previous studies of NCs assembled *in vitro* or purified from infected cells that possess weak global symmetry prior to GP interaction (Mukhopadhyay et al. 2002). We expect these deviations from global symmetry to be a feature of GP-driven budding in other spherical, enveloped viruses, with potential biological significance related to rate of budding and membrane scission, and the potential for more rapid disassembly during entry in infected cells (Lázaro, Mukhopadhyay, and Hagan 2018c).

Lateral E1 self-interactions guide assembly of the alphavirus icosahedral lattice, though the essential molecular interfaces have yet to be defined and validated(Ekström, Liljeström, and Garoff 1994). The alphavirus icosahedron is generally characterized by loose interactions between GP trimers, where the spacing is proposed to facilitate a transition of E1 fusion proteins from the pre-fusion to post-fusion conformation. In our structure, we also observed a close packing of E1 molecules from neighboring trimers at the 5-fold axis and identified nearby protein interfaces. Deletion of E1 residues D151,H152 significantly reduced MAYV growth, presumably through disruption of an interface between two quasi-3-fold spike trimers. Recently, a revertant mutation to the assembly-impaired SFV E2 H348A/H352A double mutant was mapped to an inter-trimer contact point in EI domain III at the five-fold axis, providing additional evidence this region is an important determinant of alphavirus particle formation (Byrd and Kielian, 2019). Our cryo-EM structure can be used as a guide to perform mutagenesis experiments to determine relative contributions of the interfacial residues around the five-fold axis to the stability of the virus particle.

As previously described, the outer GP shell is linked to NC in the mature virus particle by direct interaction between the Y-X-L motif of the E2 cytosolic tail and hydrophobic CP binding pocket. The correct orientation of E2 endodomain presented at the cell surface is critical for NC interaction, and it has been suggested interactions just above the viral membrane between residues in E2 subD and E1 are important for correct positioning of the CP binding site (Byrd and Kielian 2019). Within E2 subD we observe E1-E2 interactions between E2H348-E1T403 as well as E2P351-E1W407, comparable to deposited heterodimer structures of other alphaviruses (Chen et al. 2018). First noted in the cryo-EM structure of VEEV, unassigned density exists near two sequential, completely conserved tyrosines (E2-Y359,Y360) in E2subD (corresponding to MAYV E2-Y358,Y359). In our map we observed similar unassigned density near the outer leaflet of the viral envelope, situated roughly parallel and contacting E2subD in a highly lipophilic pocket above E1-E2 TM helices. Mutagenesis of conserved, sequential tyrosines within E2subD (E2-Y357,Y358,Y359) to alanine resulted in a significant decrease in MAYV production, presumably through disruption of a conserved contact between Y358 and pocket factor. This suggests the pocket factor can modulate particle assembly, presumably through stabilization of E2subD-E1 interactions and the E2 tail. More work remains to evaluate the specific role of interactions with lipophilic factor and nearby E1-E2 polar contacts, which can be essential for properly orienting both the E1 icosahedral lattice and cytosolic CP-binding motif.

In summary, deviations from global symmetry should be viewed as a general structural property of alphavirus particles and warrants further investigation in other icosahedral, enveloped virus particles. The polarity of structural order within particles gives rise to interesting hypotheses about the assembly of icosahedral enveloped viruses driven by membrane protein interactions and how it might differ from cytosolic assembly events. From the near-atomic resolution structure of MAYV it was possible to identify and validate targets for disruption of virus assembly. This includes targeting inter-trimer interactions and the lipophilic pocket factor of the E1-E2 heterodimer. We anticipate this structure will serve as a valuable resource for investigating MAYV particle assembly and pathogenicity, identifying structural conservation in other alphavirus particles, and designing future viral inhibitors.

## Materials and Methods

### Virus purification

Vero cells were prepared to 80–90% confluence and inoculated with MAYV at a multiplicity of 0.1 plaque-forming units per cell. Infected cells were incubated at 37°C for 2 days or until cytopathic effects were observed. Cellular debris was removed from the culture supernatant by centrifugation for 5–10 min at 1000–2000 *g*. Virus was concentrated by precipitation with 7% polyethylene glycol 6000 and 2.3% NaCl at 4°C for 12 h. Virus was then pelleted by centrifugation at ⍰2500 *g* for 30 min and gently resuspended in 2 ml TEN buffer (0.05 M Tris–HCl, pH 7.4, 0.1 M NaCl and 0.001 M EDTA). The virus suspension was purified by centrifugation through a 20–70% continuous sucrose/TEN gradient for 60 min at 35 000 *g*. The virus band was harvested and centrifuged 5 × through Amicon 100 kDa filter (Ultra-4 Cat. No. UFC810024), resuspending each time to maximum load volume with TEN. The purified virus was harvested in the minimal remaining volume after final centrifugation.

### Cryo-EM sample preparation and data acquisition

The preparation of MAYV particles was applied to Quantifoil copper EM grids with a holey-carbon film and plunge frozen in liquid ethane using a FEI Vitrobot Mark IV freezing apparatus. The frozen-hydrated MAYV grids were examined with a JEM 3200FSC microscope, operated at liquid-nitrogen temperature, and equipped with a Gatan K2 Summit direct-electron detector. Images were collected in super-resolution mode at 30,000x magnification and binned by 2 for processing of images at a pixel size of 1.28Å . Each image exposure was for 5 seconds, with an electron dose rate of 7 electrons/pixel/second, resulting in a dose of 35 electrons on the specimen over a total of 25 frames. In total, approximately 1100 images were collected, and 22,000 particle images were selected for further processing. Micrographs were subjected to motion correction and dose-weighting using Unblur software.

### Cryo-EM data processing

An initial model from the 22,314 selected particles was first generated with the eman2 software package (Tang et al. 2007). The particles were then processed in jspr 3d-reconstruction software, in which ctf parameters were estimated and particle centers and orientations were iteratively refined with icosahedral symmetry to give a density map at 4.7Å resolution after per-particle defocus and per-particle astigmatism corrections (Guo and Jiang 2014). Subparticles were generated by expanding symmetry from the icosahedral orientation of each particle image to all 60 ASU orientations, followed by constrained refinement of subparticles using c5 symmetry. Sorting the subparticles by phase residual agreement to a matching subvolume of the original icosahedral map increased resolution to ∼4.5Å. Non-crystallographic symmetry was imposed to average the four subunits within each asymmetric unit. This was done by loosely extracting volumes of each subunit in UCSF chimera and computing an all-vs-all alignment using EMAN2 *e2spt_hac*.*py* program. This resulted in a final density map of the subunit ectodomain at 4.2Å resolution and endodomain at 4.4Å resolution computed by gold-standard fourier shell correlation. The figures were prepared using UCSF Chimera (Pettersen et al. 2004) or UCSF Chimera X (Goddard et al. 2018).

An asymmetric reconstruction of MAYV using the 22,000 selected particles was performed using RELION 3.0 (Fernandez-Leiro and Scheres, n.d.; Scheres 2012). Prior to refinement, micrographs were further binned to a pixel size of 2Å, motion-corrected using MotionCor2, and ctf parameters were estimated using CTFFIND4 (S. Q. Zheng et al. 2017; Rohou and Grigorieff 2015). An icosahedrally-symmetric starting model of MAYV was low-pass filtered to 60Å to blur the high resolution features. Orientations of each particle were determined without imposing symmetry, with global angular search value of 3.7° and local search angular increment of 0.9°. The orientation search converged after 26 iterations, resulting in an asymmetric reconstruction at ∼9Å resolution using gold standard FSC criteria.

### Model building and refinement

For proteins E1, E2, E3 and CP, homology models were first generated from the MAYV primary sequence using Phyre protein structure prediction server(Kelley et al. 2018). These models were fit into the respective density from the single MAYV protomer after NCS averaging using UCSF Chimera and refined in real space using Phenix software taking into account stereochemical and secondary structure restraints(Kelley et al. 2018; Pettersen et al. 2004). Further, an iterative process of manual editing of the polypeptide backbone and rotamer placements using COOT and further Phenix real space refinements was performed until the refinement statistics and visual fit of atoms showed no clear improvement. To assess the resolvability of the map and the quality of the model, Q-scores were calculated for each residue with the final model (Pintilie et al. 2020a). Refinement statistics of the structural model are listed in Appendix Table 1, while plots of the Q-score values for each protein can be found in Figures EV 2 and 3.

### Reporter virus development

The infectious clone of wild type MAYV CH strain was obtained from the World Reference Center for Emerging Viruses and Arboviruses (WRCEVA) at the University of Texas Medical Branch, Galveston, TX. The nanoluciferase gene was inserted between E3 and E2 after the furin cleavage site within the MAYV genome using standard overlapping PCR approach to generate a MAYV-NLuc construct. Site-directed mutagenesis was performed by standard overlapping PCR approach. Molecular clones of mutant viruses were confirmed by sanger sequencing prior to rescue.

### Assessing viral replication

Viral genomic RNA from the wild type and mutant MAYV-NLuc clones were produced by linearization with PacI then *in vitro* transcribed from an SP6 promoter using the mMESSAGE mMACHINE kit (ThermoFisher Scientific). 10 Lg of RNA was electroporated into 1×10^7^ BHK-21 cells which were grown in 15 ml DMEM media. 0.5 ml of culture supernatants were collected every 6 hours and nanoluciferase activities associated with released virus in the supernatants were measured using Nano-Glo luciferase assay system (Promega).

## Supporting information

Expanded View Figures

## Acknowledgements

We thank Joanita Jakana and all staff at the National Center for Macromolecular Imaging (NCMI) for assistance with data collections, general support, and expert maintenance of the cryo-EM facilities. This work was supported by grants from the National Institute of Allergy and Infectious Diseases of the National Institutes of Health under Award Numbers P41GM103832 and P01AI120943 to WC, K22AI125474 and R01AI153433 to AJA, and R24AI120942 to SCW.

## Author Contributions

W.C., A.A., and S.W. supervised the study. A.A. prepared the sample for cryo-EM. J.J. prepared virus mutant experiments. D.C. and J.K. performed cryo-EM sample preparation. J.K. and D.C. collected cryo-EM data. D.C. and J.K. performed cryo-EM image processing and structure determination; J.K. built and refined the model; D.C., J.K., A.A., J.J., S.W., and W.C. analyzed data. D.C. prepared figures. D.C. and W.C. wrote the manuscript with input from all other authors.

## Data Deposition

Cryo-EM maps of the Mayaro virus E1-E2-CP subunit with its associated atomic model have been deposited in the wwPDB OneDep System under EMD accession code EMD-XXXXX and PDB ID code XXXX.

## Expanded View Figure Legends

**EV Figure 1**. Single particle cryo-EM data processing of MAYV virion. (a) Workflow diagram of data processing, including gold-standard FSC curves of (I) subparticle focused region and (II) after NCS averaging of four quasi-equivalent subunits. (b) Representative density of E1/E2 ectodomain (top) and endodomain (bottom) after (I) initial refinement using icosahedral symmetry, (II) focused sub-particle refinement, and (III) focused sub-particle refinement + NCS averaging.

**EV Figure 2**. Model validation of the E1/E2 heterodimer ectodomains. (a) Transparent density map with backbone model colored by residue Q-score for E2 (top) and E1 (bottom) ectodomains (b). Q-score for each amino acid residue in E2 (top) and E1 (bottom) model and 4.2Å map. Red line (0.41) represents expected Q value at 4Å resolution based on correlation between Q-scores and map resolution. (c) Sample density of E2 (top) including high- and low-scoring stretches of residue Q-scores and E1 (bottom), with residue (black) and Q-score (blue) labels.

**EV Figure 3**. Model validation of the MAYV endodomain. (a) Transparent density map of endodomain with backbone model colored by residue Q-score. (b) Q-score for each residue in capsid protein (CP) and 4.4Å map, red line (0.41) represents expected residue Q-score in 4Å map. (c) Sample density of CP with model residues (black) and Q-scores (blue) indicated. (d) Interaction interface between E2 (green) and CP (cyan). Amino acid residues identified as forming E2-CP contacts are displayed with zoned density. Table lists all residues identified in the forming E2-CP interface.

**EV Figure 4**. Comparison of alphavirus structures. Overlay of our atomic MAYV model with other deposited alphavirus atomic models, aligned and colored by rmsd.

**EV Figure 5**. Glycosylation sites of MAYV. (a) Cryo-EM map of the MAYV trimer with glycans shown (green). (b) E1/E2 heterodimer density with zoom-in views of both N-linked glycans.

## Appendix Figure Legends

**Appendix Figure 1**. Sequence alignment of alphavirus E1, E2, CP. Alignment performed using NPS and ClustalW, formatted with ESPript.

**Appendix Movie 1**. Pocket factor location and interactions. Related to Figure 5. Unmodeled density in the MAYV E1/E2/E3/CP subunit, initially colored as grey, sits in a lipophilic pocket between E1 and E2. Residues of E1 and E2 that form continuous density with this unassigned factor are displayed.

**Appendix Table S1**. Cryo-EM data collection, refinement and validation statistics.

