## Supplementary material for "Near-atomic resolution Cryo-EM structure of Mayaro virus identifies key structural determinants of alphavirus particle formation": Expanded View Figures

**EV Figure 1**

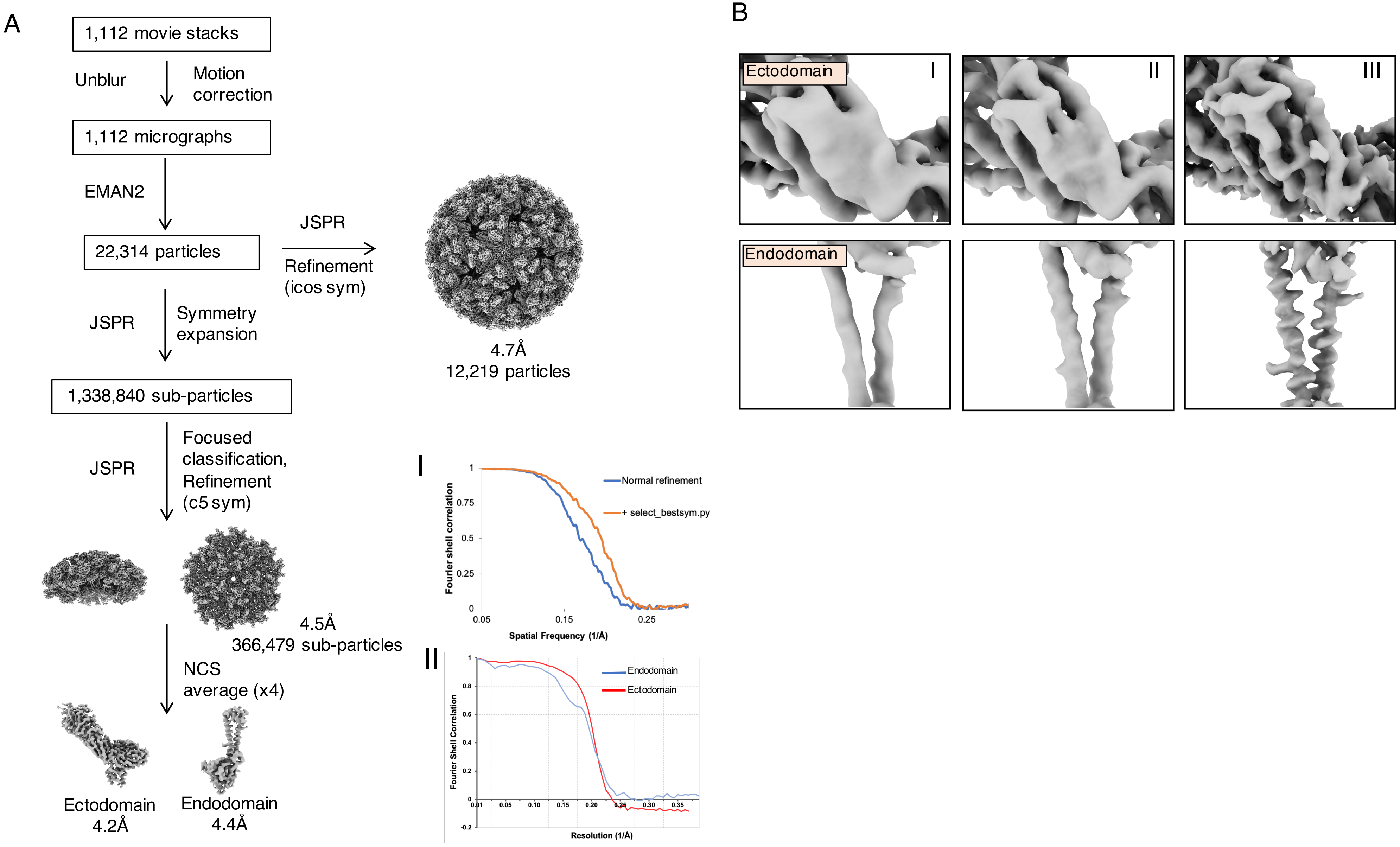

A
1,112 movie stacks
Unblur
Motion correction
1,112 micrographs
EMAN2
22,314 particles
JSPR
Refinement (icos sym)
4.7Å
12,219 particles
JSPR
Symmetry expansion
1,338,840 sub-particles
JSPR
Focused classification, Refinement (c5 sym)
4.5Å
366,479 sub-particles
NCS average (x4)
Ectodomain
4.2Å
Endodomain
4.4Å
I
Fourier shell correlation
Spatial Frequency (1/Å)
Normal refinement
+ select_bestsym.py
II
Fourier Shell Correlation
Resolution (1/Å)
Endodomain
Ectodomain
B
Ectodomain
Endodomain
I
II
III

### EV Figure 2

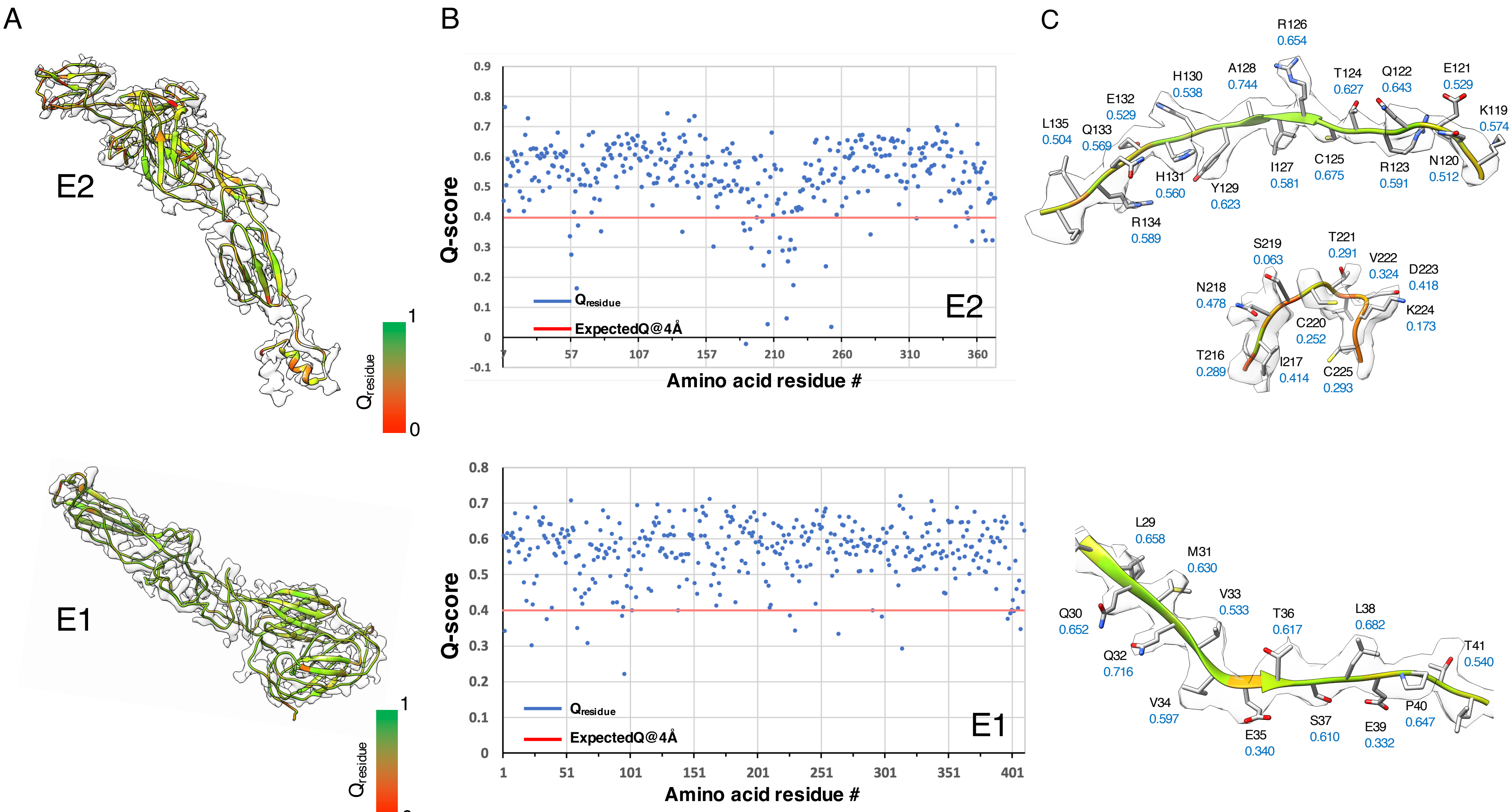
A
E2
Qresidue
1
0
E1
Qresidue
1
0
B
Q-score
0.9
0.8
0.7
0.6
0.5
0.4
0.3
0.2
0.1
0
-0.1
7
57
107
157
210
260
310
360
Amino acid residue #
Qresidue
ExpectedQ@4Å
E2
Q-score
0.8
0.7
0.6
0.5
0.4
0.3
0.2
0.1
0
1
51
101
151
201
251
301
351
401
Amino acid residue #
Qresidue
ExpectedQ@4Å
E1
C
R126 0.654
H130 0.538
A128 0.744
T124 0.627
Q122 0.643
E121 0.529
E132 0.529
K119 0.574
L135 0.504
Q133 0.569
H131 0.560
Y129 0.623
I127 0.581
C125 0.675
R123 0.591
N120 0.512
R134 0.589
S219 0.063
T221 0.291
V222 0.324
D223 0.418
N218 0.478
K224 0.173
C220 0.252
T216 0.289
I217 0.414
C225 0.293
L29 0.658
M31 0.630
V33 0.533
Q30 0.652
T36 0.617
L38 0.682
T41 0.540
Q32 0.716
V34 0.597
E35 0.340
S37 0.610
E39 0.332
P40 0.647

**EV Figure 3**

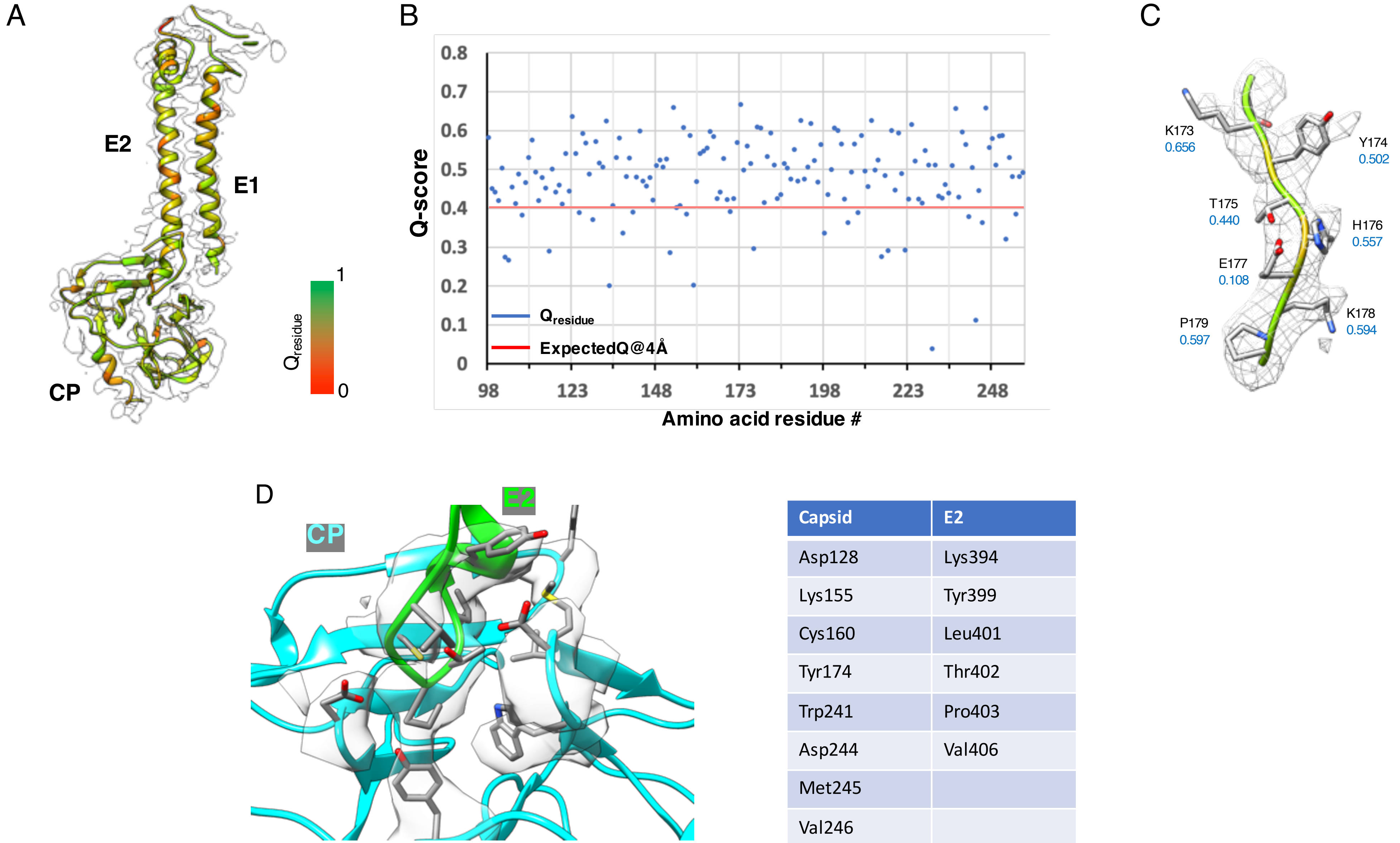


| Capsid | E2 |
|---|---|
| Asp128 | Lys394 |
| Lys155 | Tyr399 |
| Cys160 | Leu401 |
| Tyr174 | Thr402 |
| Trp241 | Pro403 |
| Asp244 | Val406 |
| Met245 | |
| Val246 | |

### EV Figure 4

#### E2 ectodomain

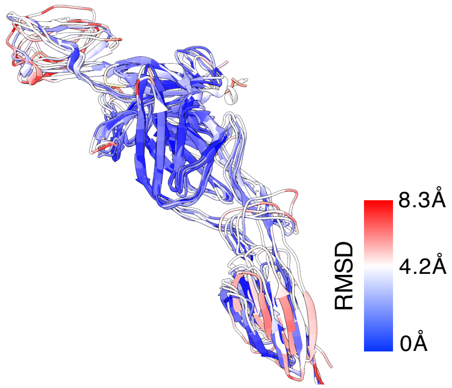


| Model | RMSD |
|---|---|
| MAYV | reference |
| CHIKV (PDB:3n41) | 2.992 |
| SINV (PDB:6imm) | 1.953Å |
| EEEV (PDB:6mx4) | 3.753Å |
| VEEV (PDB:3joc) | 2.970Å |

#### E2 endodomain

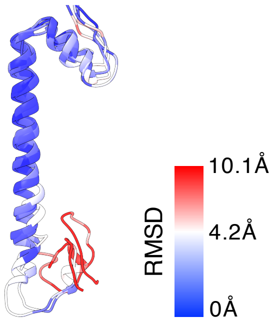


| Model | RMSD |
|---|---|
| MAYV | reference |
| SINV (PDB:6imm) | 1.316Å |
| EEEV (PDB:6mx4) | 5.331Å |
| VEEV (PDB:3joc) | 3.479Å |

#### E1 ectodomain

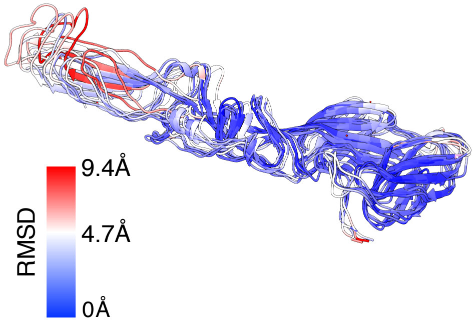


| Model | RMSD |
|---|---|
| MAYV | reference |
| CHIKV (PDB: | 2.861Å |
| SINV (PDB:6imm) | 2.351Å |
| EEEV (PDB:6mx4) | 4.739Å |
| VEEV (PDB:3joc) | 2.678Å |

#### E1 endodomain

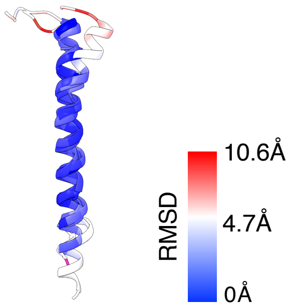


| Model | RMSD |
|---|---|
| MAYV | reference |
| SINV (PDB:6imm) | 1.501Å |
| EEEV (PDB:6mx4) | 2.011Å |
| VEEV (PDB:3joc) | 2.819Å |

#### Capsid

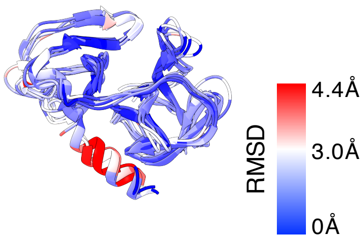


| Model | RMSD |
|---|---|
| MAYV | reference |
| SINV (PDB:1svp) | 2.591Å |
| CHIKV (PDB:5h23) | 1.422Å |
| EEEV (PDB:6mx4) | 2.034Å |
| VEEV (PDB:3joc) | 1.602Å |

### EV Figure 5

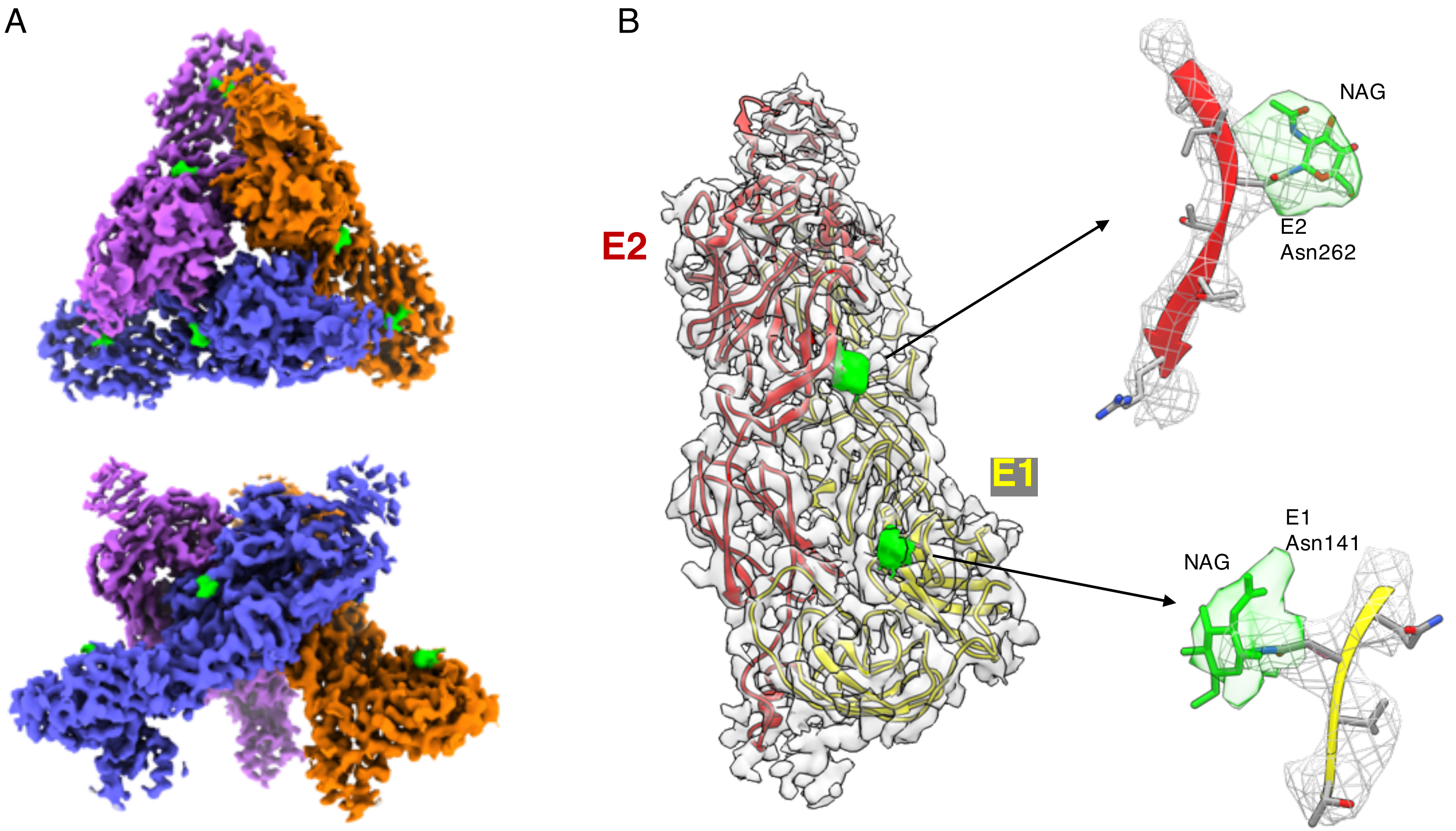
