## Appendix Table 1 for "Near-atomic resolution Cryo-EM structure of Mayaro virus identifies key structural determinants of alphavirus particle formation"

Table S1. Cryo-EM data collection, refinement, and validation statistics.

MAYV subunit

(EMDB-XXXXX)

(PDB ID: XXXX)

**Data collection and processing**

Magnification	28k

Voltage (kV)	300

Electron exposure (e^-^/Å^2^)	35

Defocus range (um)	1.2-3.5

Pixel size (Å)	1.28

Initial particle images (no.)	22,314

Final particle images (no.)	12,219

Symmetry imposed	Icos

Map resolution (Å)	4.7

FSC threshold	0.143

Symmetry expansion data processing

Initial subparticle images (no)	1,338,840

Final subparticle images (no.)	366,479

Symmetry imposed	C5

Map resolution (Å)	4.4

FSC threshold	0.143

**Refinement**

Map sharpening *B* factor (Å^2^)	-240

Model composition

Non-hydrogen atoms	7714

Protein residues	1002

R.M.S. deviations

Bond lengths (Å)	0.0093

Bond angles (°)	1.47

Validation

MolProbity score	2.35

Clashscore	7.91

Poor rotamers (%)	2

Ramachandran plot

Favored (%)	87.5

Allowed (%)	12.1

Disallowed (%)	0.4
